# Observation of general and specific adverse effects of organic solvents on *Caenorhabditis elegans* by toxicity testing using behavioral analysis

**DOI:** 10.1101/2024.09.21.614284

**Authors:** Masahiro Tomioka

## Abstract

Novel chemical compounds are continuously being developed and are used in various industries as well as in our daily lives. Workers in industries estimate and avoid chemical hazardous risks based on published information on chemical toxicities. However, it is challenging to avoid hazards caused by chemicals with unknown toxicity. Therefore, it is necessary to understand the toxicities of chemicals in high-throughput and multifaceted ways. In this study, I develop a high-throughput method to assess chemical toxicities through the quantitative measurement of behavior in *Caenorhabditis elegans*. I determine the acute toxicity of 30 organic solvents that are widely used in industries with motility as an endpoint. The adverse effects of organic solvents on motility were proportional to the lipid solubility of chemicals, which is similar to the positive relationship between the anesthetic effects of volatile organic chemicals and their lipid solubility in organisms, including humans. In addition to the general toxicity on motility, organic solvents caused adverse effects on posture during locomotion in different ways depending on the chemical’s functional group. Alcohols and cellosolves, both of which have a hydroxyl group, reduced the amplitude of body bending, whereas ketones and acetate esters, both of which have a ketone group, increased it during undulatory locomotion. Furthermore, organic solvents caused defects in behavioral plasticity based on the association of starvation and chemical signals at concentrations lower than those that affect locomotion, which suggests that sensory processing is more susceptible to organic solvents than motor function. This study provides a high-throughput method for acute chemical toxicity testing and new insights into the behavioral toxicity of organic solvents based on comprehensive behavioral analysis.

## Introduction

Organic solvents are used in various industries as well as in our daily lives. Many organic solvents are amphiphilic and can penetrate the epithelium and thus have a risk of being a health hazard to organisms, especially workers in industries. Commonly used organic solvents, such as alcohol, ketone, and ether, can cross the blood-brain barrier and cause neurotoxicity when exposed to high levels of these chemicals. For example, low concentrations of benzyl alcohol are generally used in daily life, such as in foods, drugs, and cosmetics. Meanwhile, benzyl alcohol intoxication has been reported in workers using paint strippers containing high concentrations of benzyl alcohol [1, 2]. The intoxicated patients collapsed during paint stripping in a state of unconsciousness, and severe damage to the central nervous system has been reported [1]. To reduce the hazardous chemical risks, a hazard assessment has been performed using the information on chemical toxicities, such as the GHS (Globally Harmonized System of Classification and Labelling of Chemicals) [3, 4], which classifies them based on accident cases and toxicity studies on animals. However, there are many chemicals with unknown hazards. Benzyl alcohol had not been classified as a chemical with specific target organ toxicity to the brain before serious accidents occurred in Japan. To predict chemical hazards and prevent accidents before they happen, it is important to fundamentally understand the toxic effects of chemicals and the mechanisms of the adverse effects on organisms.

Chemical toxicities occur by penetration of the chemicals into cells and their subsequent interactions with biological molecules. Cell permeability depends on the chemical’s physicochemical characteristics, including molecular size, melting point, and lipid solubility. The externally incorporated chemicals interact with biological molecules, such as proteins, lipids, and DNA, to exert their toxic effects. Exposure to high levels of organic solvents causes acute toxicity to the brain, followed by disturbance of consciousness similar to exposure to volatile anesthetics. More than 120 years ago, Meyer and Overton independently reported that the anesthetic effects of volatile organic chemicals increase with increasing lipid solubility of the chemicals [5, 6]. Subsequent studies have shown that increased lipid solubility elevates the passive transport into cells through the plasma membrane, which is constituted of a lipid bilayer, and could affect the function of multiple lipids and proteins within the cell membrane, such as ion channels and synaptic proteins, although the mechanisms are not fully understood [7]. These notions explain, at least in part, the Meyer–Overton hypothesis, which is the positive relationship between the anesthetic effect and lipid solubility of organic chemicals. Similarly, it was reported that lipid solubility was linearly correlated with the adverse effects on growth in microorganisms, such as yeast [8]. Thus, lipid solubility increases the general toxicity of organic chemicals in multiple cell types across species.

The nematode *Caenorhabditis elegans* has been widely used as a powerful model organism in basic science research, such as genetics, molecular biology, and neurobiology. Moreover, its use in toxicology research has increased in recent years [9]. Several studies have shown similarities between *C. elegans* and higher organisms in the toxicity ranking of chemicals [10]. Although the site of action of toxicants in the *C. elegans* body has not been extensively studied, biological targets appear to be relatively similar between *C. elegans* and higher organisms based on the conservation of genes that compose cellular structure, molecular signaling, and enzymes that produce biological materials [9]. A variety of biological analyses for molecular biology, cell biology, and phenotype observation, including development, reproduction, and behavior, have been established. Furthermore, a small body size of approximately 1 mm in adults and a fast life cycle of approximately 3 days from egg to mature adult under optimal conditions enable high-throughput screening, such as chemical screening, using large quantities of the nematodes [11]. Therefore, *C. elegans* is an ideal animal model to understand chemical toxicities in organisms and their underlying cellular and molecular mechanisms. Although a standardized protocol for assessing environmental toxicity has been developed [12], no standardized protocol exists for determining chemical toxicity in humans using *C. elegans*.

In this study, a simple method for evaluating the acute behavioral toxicity of chemicals using *C. elegans* was developed, and 30 organic solvents were selected that are commonly used in chemical industries for the initial study. Adverse effects on locomotion were quantified using a tracking system after exposure of the nematodes to organic solvents on agar plates. The motility of nematodes substantially decreased and eventually ceased after exposure to any of the 30 organic solvents. The paralyzed nematodes recovered after incubation without the organic solvents, which suggests that the organic solvents have an anesthetic effect on the nematodes. The extent of the anesthetic effects was proportional to the octanol–water partition coefficient, which reflects the lipid solubility. Therefore, the Meyer–Overton relationship was observed in the adverse effects of the organic solvents, which is consistent with the previous reports demonstrating the positive relationship between lipid solubility and the potency of several anesthetics in *C. elegans* [13].

Furthermore, the specific effects of organic solvents on the locomotion patterns in which the phenotypes were found to qualitatively differ depending on the organic solvent’s functional group. It was reported that the amplitude of the body bends during undulatory locomotion was regulated by motor neurons that regulate the head angle, such as SMB cholinergic motor neurons and RME GABAergic motor neurons [14, 15]. The amplitude of the body bends during undulatory locomotion was quantified, and the adverse effects on body bending were observed after exposure to organic solvents at concentrations lower than those that resulted in complete paralysis. The amplitude of the body bends considerably increased after exposure to chemicals with short-chain ketone and acetate ester groups. Conversely, it decreased after exposure to chemicals with short-chain alcohol and cellosolve groups. These findings imply that organic solvents affect the neural circuit that regulates body bending in different ways depending on the solvent’s distinct functional groups.

Finally, it was demonstrated that organic solvents caused substantial defects in salt chemotaxis learning, a paradigm of learned behavior in salt chemotaxis, at concentrations lower than those that affected locomotion [16]. These adverse effects were dependent on glutamatergic signaling that regulates sensory processing, including salt chemotaxis learning, which implies that low concentrations of organic solvents cause adverse effects on the sensory neural circuits. This study suggests that organic solvents cause adverse effects on multiple systems, including motor and sensory neural circuities, in a concentration-dependent manner in *C. elegans*.

## Materials and Methods

### Cultivation of *C. elegans*

The nematode *C. elegans* was cultivated on nematode growth medium (NGM) plates using a standard protocol [17]. The wild type N2, the *unc-29(e193)* CB193, the *eat-4(ky5)* MT6308, and the *daf-2c(pe2722)* JN2722 *C. elegans* strains were used. The *Escherichia coli* OP50 bacterial strain was used as a food source. To prepare synchronized populations of *C. elegans* for locomotion analysis, eggs were harvested from gravid adult nematodes by treatment with alkaline hypochlorite solution, and approximately 600 eggs were cultivated on NGM plates at 18°C–20°C for 4–5 days until they reached the young adult stage.

### Recording of the nematodes’ movement after organic solvent exposure

The synchronized populations of adult *C. elegans* were transferred from growth plates to 500 µL buffer solution (5 mM potassium phosphate, 1 mM CaCl_2_, 1 mM MgSO_4,_ and 0.05% gelatin), which included the organic solvents. To determine the MC_<50_ values, organic solvents were dissolved in buffer at 4 to 7 different concentrations within a range of 0.125% to 6%, and the nematodes were exposed for 15, 30, 60, or 120 min. For the control group, a buffer solution without organic solvents was used for the exposure procedure. To examine recovery from paralysis, the nematodes were soaked in a buffer including organic solvents at their minimum concentration that caused complete paralysis, which is defined as the average locomotion speed below 10 µm/s for 2 h, unless otherwise stated. The nematodes were then transferred into a buffer without organic solvent and incubated for 30, 60, or 120 min. After this soaking procedure, a 30 µL droplet with approximately 50 nematodes was transferred onto a test plate (5 mM potassium phosphate, 1 mM CaCl_2_, 1 mM MgSO_4,_ and 2% Bacto agar), and excess liquid was removed with paper wipers. The nematodes’ movement on the test plates was captured under a stereo microscope (S6E, Leica Microsystems, Germany) with a USB CMOS camera (J-scope, Sato Shoji Corporation, Japan) at 30 frames per second for approximately 1 min.

### Tracking analysis of *C. elegans* locomotion

*C. elegans* locomotion was analyzed using the WormLab software (MBF Bioscience, Vermont, USA) with 1500 frame image sequences (50 s movie at 30 frames per second). The center point of each nematode on the agar plates was tracked (Figure 1A; Supporting Movie 1), and the movement speed of each trace was determined, where the plus value represents forward movement, the minus value represents reverse movement, and the absolute value below 30 µm/s was defined as a pause (Figure 1B). The locomotion speed of each trace was determined as the average speed during forward movement, and the mean locomotion speed of all traces was plotted along with the standard error of the mean (SEM) on a graph after normalization with a control (Figure 1D, right). Immobility was determined as a ratio of a pause in each trace, and the mean value of all traces along with the SEM was plotted on a graph (Figure 1D, left). The length of each nematode was determined as the length from head to tail along the central axis, and the mean length of all nematodes was plotted along with the SEM on a graph (Figure 3A). The amplitude of the body bends during locomotion was determined as the average centroid displacement, which is the distance between the midpoint and the average location of the central axis points of nematodes (Figure 5B), and the mean amplitude of each trace was plotted on violin plots (Figure 5C). At least three traces were used for the analyses.

**Figure 1.**
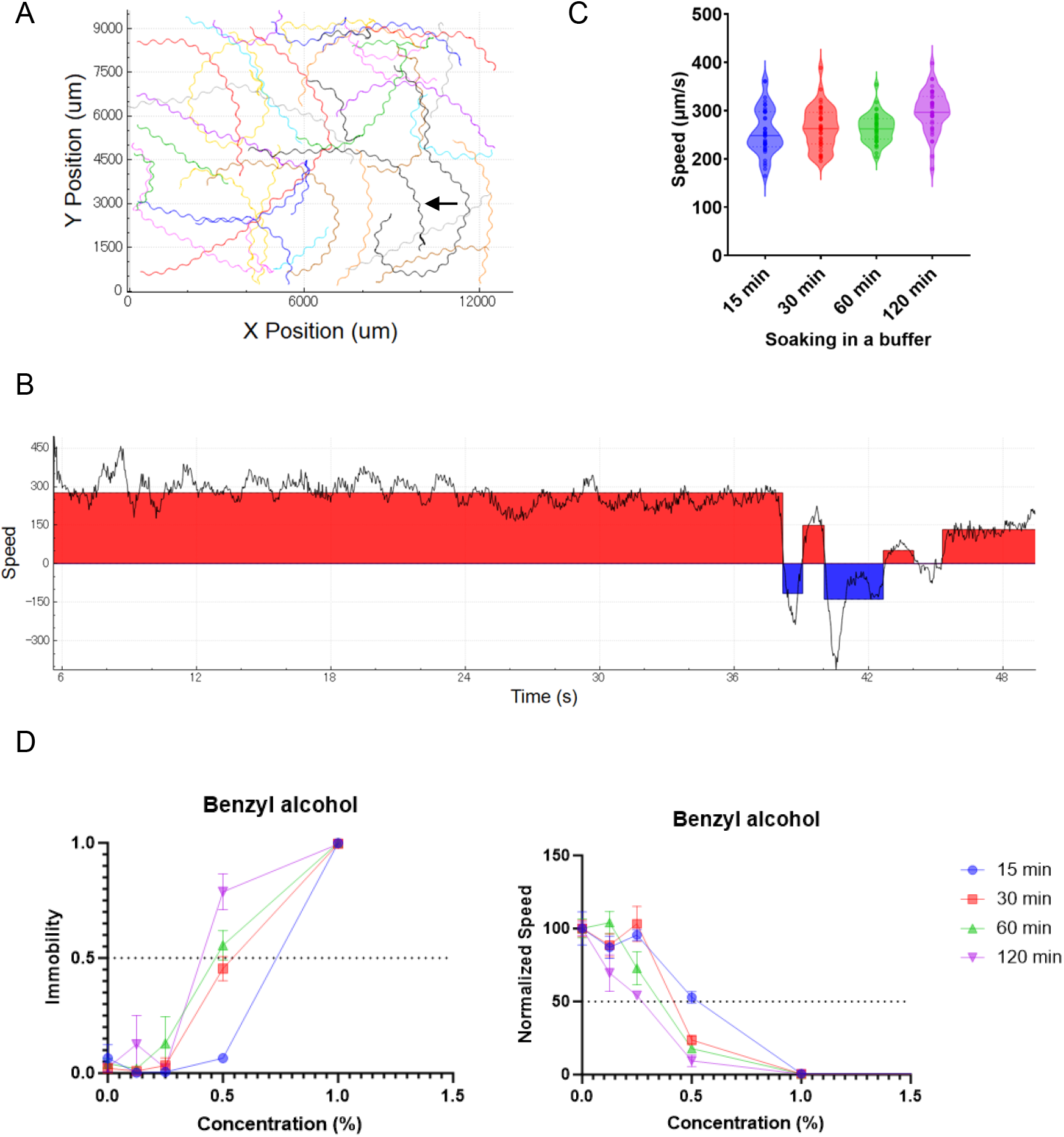
Quantification of the locomotion of *C. elegans* exploring on agar plates. (A) Representative trajectories of the nematode *C. elegans* moving on an agar plate for 50 s. A trace depicted by an arrow is used for the representative analysis of the locomotion speed shown in B. (B) Locomotion speed (µm/s) during 50 s tracking of *C. elegans*. Plus values (red region) or minus values (blue region) are defined as “Forward” or “Reverse” movements, respectively. Absolute values below 30 µm/s (colorless region) are defined as “Pause”. (C) Violin plots of the locomotion speeds after soaking in buffers without organic solvents. Each dot represents the mean locomotion speed in each trace. Solid horizontal lines represent the median. Dotted horizontal lines represent the 25^th^ and 75^th^ percentiles. (D) Immobility rate (left) and locomotion speed (right) after exposure to benzyl alcohol for 15, 30, 60, or 120 min. Each data point represents the mean ± the standard error of the mean (SEM). Locomotion speed was normalized to the average value after exposure without benzyl alcohol.

**Figure 2.**
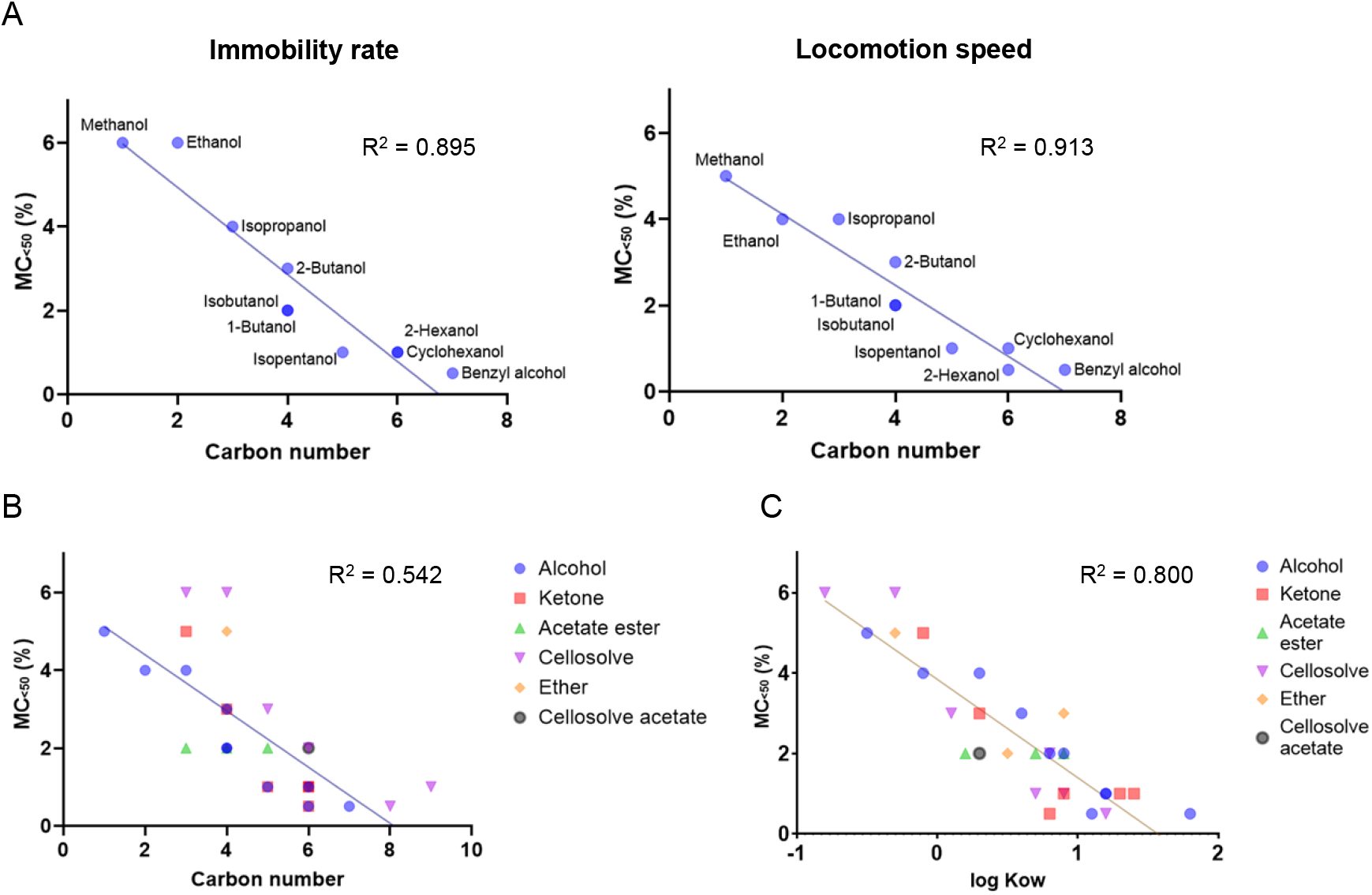
Correlation of acute behavioral toxicity with the organic solvent’s carbon number and lipid solubility. (A) Relationship between the behavioral toxicity (MC_<50_) after alcohol exposure for 1 h and the carbon number of the alcohols. The immobility rate (left) and locomotion speed (right) are used as the endpoints. (B, C) Relationship between the MC_<50_ after exposure for 1 h to organic solvents and the chemical’s carbon numbers (B) or octanol–water partition coefficients (C). Locomotion speed is used as an endpoint. R^2^ values were determined based on simple linear regression analysis.

**Figure 3.**
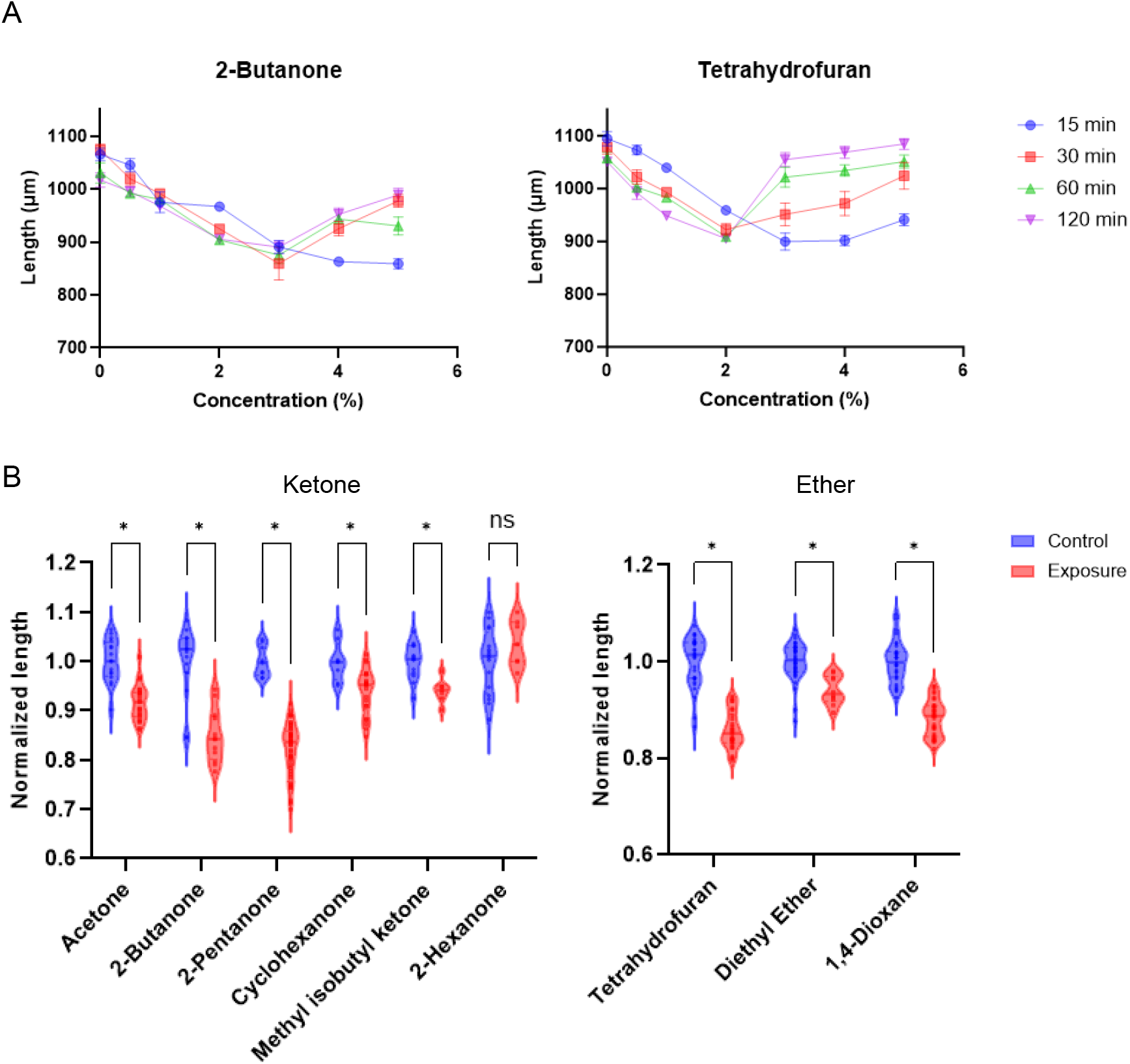
Ketone and ether exposure causes body shrinkage followed by relaxation. (A) Changes in body length after exposure to 2-butanone (left) or tetrahydrofuran (right). Each data point represents the mean ± the standard error of the mean (SEM). (B) Violin plots of the nematode’s body lengths after exposure to ketones (left) or ethers (right). The exposure concentrations are the MC_<50_ values determined by the locomotion speed after exposure for 1 h. Data were normalized to the average values of the control. Each dot represents the body length of a nematode after exposure (red) or without exposure (blue). \**P* < 0.05, unpaired t-test with Holm–Sidak correction.

**Figure 4.**
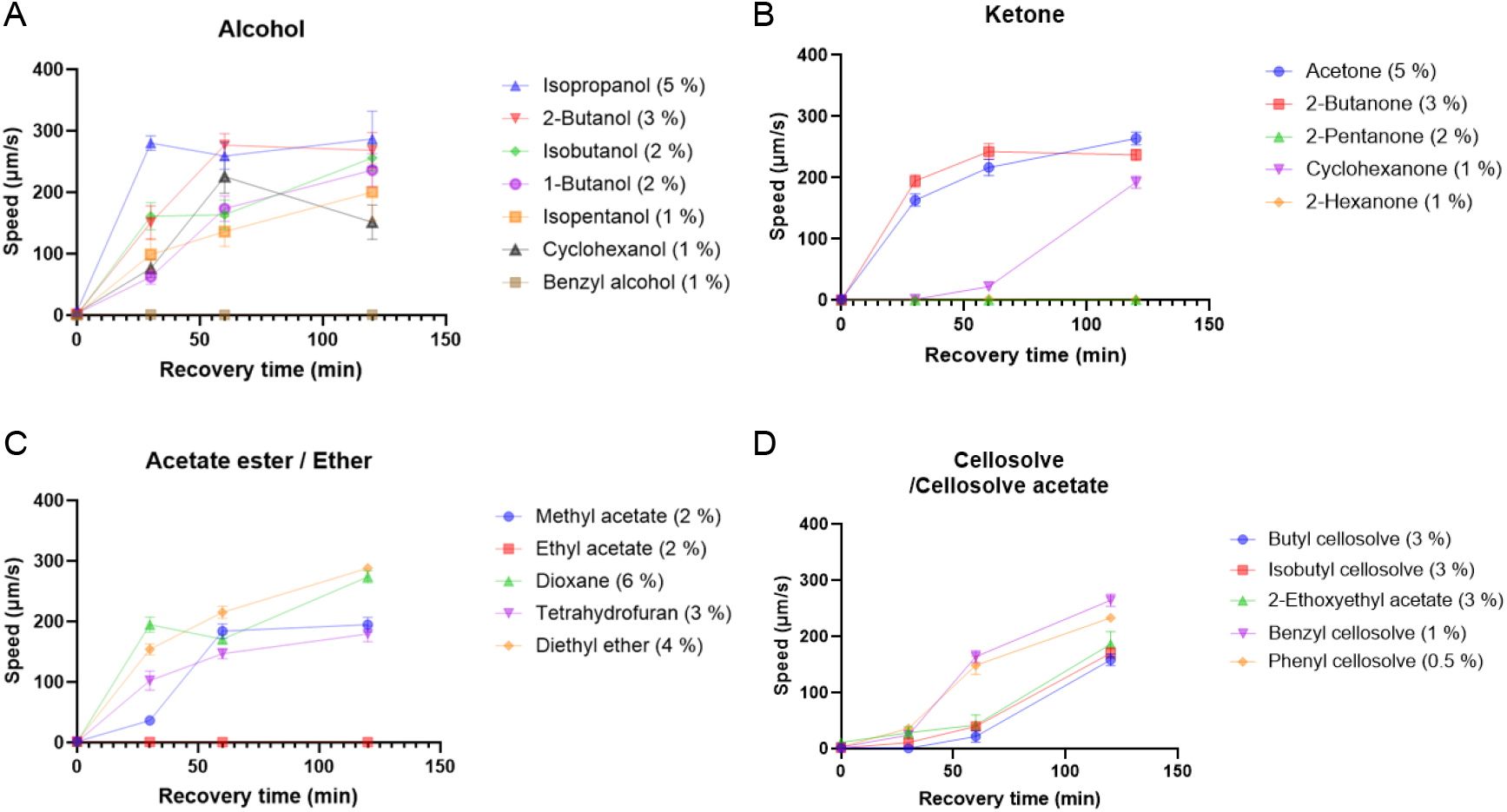
Time course of recovery from paralysis after organic solvent exposure. (A–D) Locomotion speed after recovery in a buffer after organic solvent exposure for 2 h is shown. The exposure concentrations of the organic solvents are the minimum concentrations that cause complete paralysis. Each data point represents the mean ± the standard error of the mean (SEM).

**Figure 5.**
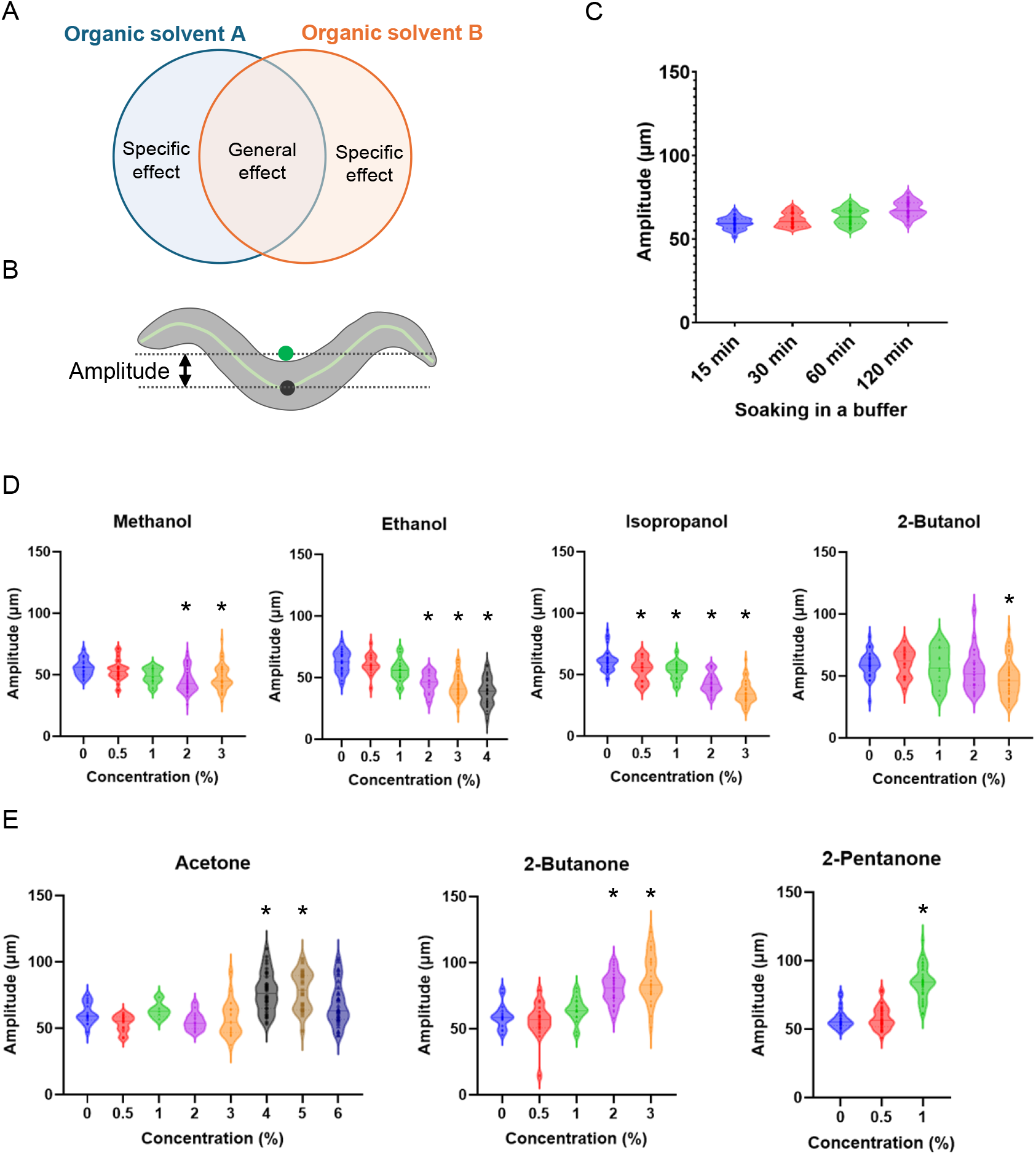
Alcohols and ketones cause adverse effects on the posture during locomotion in opposite directions. (A) Working model for the toxicity of organic solvents. Organic solvents have general toxicities that induce similar effects on an organism as well as specific toxic effects that are qualitatively different among chemicals. (B) Quantification of the amplitude of body bending during locomotion. The midpoint (black dot) and the average location of the central axis points (green dot) were shown. (C) Violin plots of amplitudes after soaking in a buffer for 15, 30, 60, or 120 min. (D, E) Violin plots of the amplitudes after soaking in a buffer containing alcohols (D) or ketones (E) for 15 min. Each dot represents the mean amplitude during each track. \**P* < 0.05, one-way ANOVA with Dunnett test, compared with the no-exposure control.

### Salt chemotaxis learning

A salt chemotaxis learning assay was performed according to a previously published procedure with some modifications [16] (Supporting Figure 14). To prepare a chemotaxis test plate, an agar block including NaCl (50 mM NaCl, 5 mM potassium phosphate, 1 mM CaCl_2_, 1 mM MgSO_4,_ and 2% Bacto agar) was placed at the edge of an agar plate (5 mM potassium phosphate, 1 mM CaCl_2_, 1 mM MgSO_4_, and 2% Bacto agar) to form a NaCl-concentration gradient by over-night incubation at room temperature (Supporting Figure 14A, right). For conditioning, the nematodes were transferred into 500 µL buffer solution (5 mM potassium phosphate, 1 mM CaCl_2_, 1 mM MgSO_4,_ and 0.05% gelatin) with or without 20 mM NaCl and incubated for 1 h, and named NaCl(+) or NaCl(−) conditioning, respectively. To assess the effect of organic solvent exposure for 1 h, the organic solvents were dissolved in a buffer solution for NaCl(+) or NaCl(−) conditioning. To assess the effect of organic solvent exposure for 1 min, nematodes were soaked in buffer solutions with or without NaCl for 1 h, then were transferred into NaCl(+) or NaCl(−) buffer, respectively, which included the organic solvents, and were incubated for 1 min. After conditioning, a droplet with approximately 50–100 nematodes was transferred onto the chemotaxis test plate, and excess liquid was removed with paper wipes. Just before transferring the nematodes onto the chemotaxis test plate, the agar block was removed, and 0.5 µL of 1 M sodium azide was spotted at both positions where the agar block was placed and the opposite side to anesthetize nematodes at those positions (Supporting Figure 14B). After the nematodes freely explored the test plate for 15 min, the numbers of anesthetized nematodes at each position were counted and the chemotaxis index was calculated, as shown in Supporting Figure 14B. When the chemotaxis index value is 1.0, all nematodes that migrated from the starting area moved toward the high-NaCl area. When the chemotaxis index value is −1.0, all nematodes that migrated from the starting area moved toward the low-NaCl area. Assays were repeated six times.

### Statistics and raw data

Statistical analyses were performed using GraphPad Prism 10. Simple linear regression analysis was performed to calculate the values of the coefficient of determination, R^2^ (Figure 2, Supporting Figure 4–6). For multiple comparison tests, an unpaired *t*-test with Holm–Sidak correction (Figure 3B, Supporting Figure 9), a one-way ANOVA with Dunnett test (Figure 5D, E, Supporting Figure 13), and a two-way ANOVA with Dunnett test (Figure 6) were used. Raw data and raw statistics are shown in Supporting Files 1–4.

**Figure 6.**
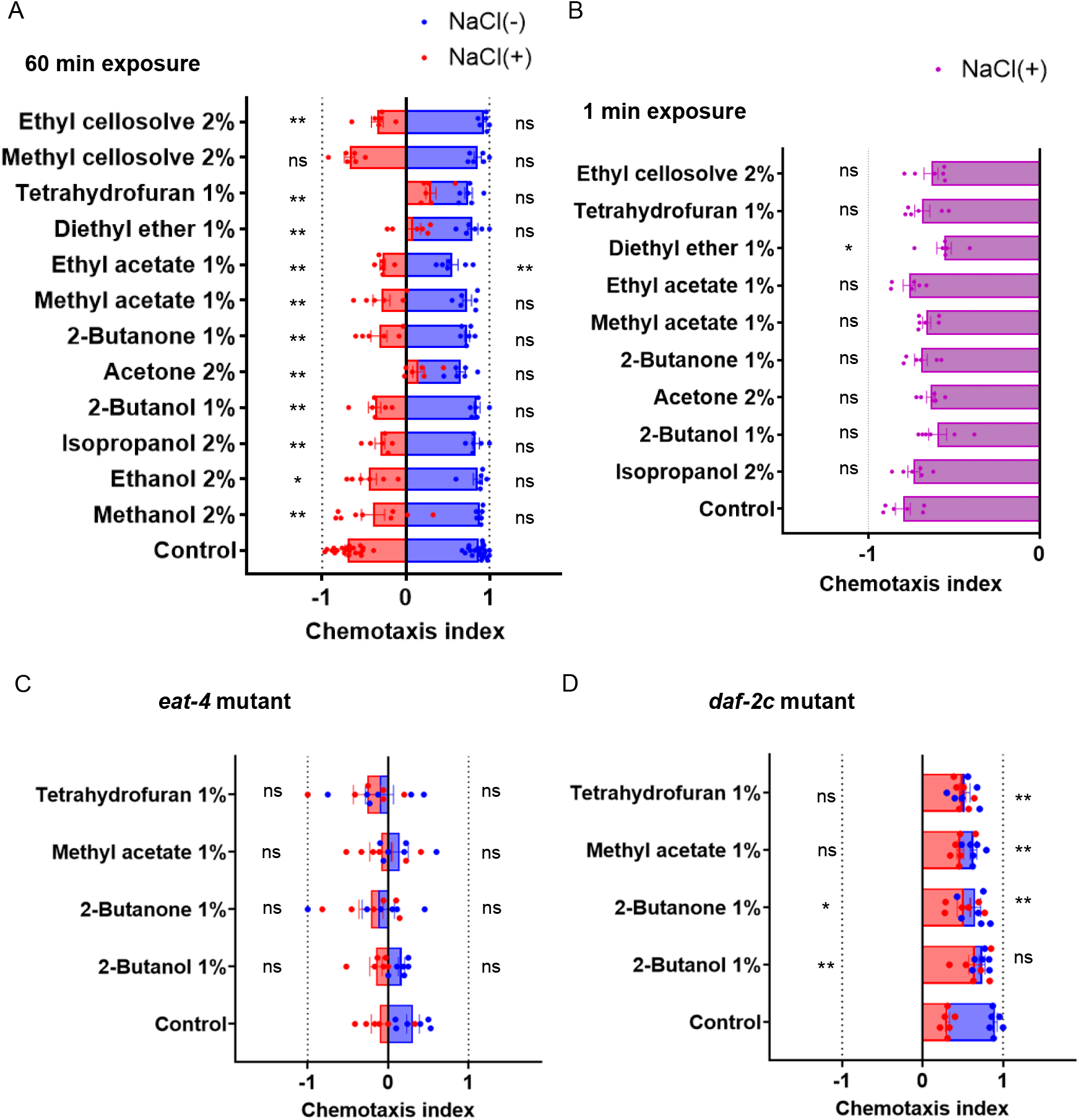
Organic solvent exposure causes defects in salt chemotaxis learning. Salt chemotaxis after soaking in buffer with or without NaCl. Nematodes were exposed to organic solvents for 1 h during the soaking procedure (A, C, D) or for 1 min after the soaking procedure (B). The exposure concentrations of organic solvents are the maximum concentrations at which no significant defect is observed in the locomotion speed after exposure for 1 h. The wild type (A, B), the *eat-4(ky5)* mutant (C), and the *daf-2c(pe2722)* mutant (D) were used. The chemotaxis index represents the extent and direction of chemotaxis, where +1 and −1 indicate that all nematodes are attracted to or avoid NaCl, respectively. Each dot represents the chemotaxis index calculated for each trial. n = 6 assays. The bar and error bars indicate the mean ± the standard error of the mean (SEM). \**P* < 0.05; \*\**P* < 0.01, two-way ANOVA with Dunnett test, compared with the no-exposure control.

## Results

### Behavioral toxicity of monohydric alcohol is proportional to the carbon number of the alcohol

The locomotion of *C. elegans* on an agar plate was quantified using a tracking system of the nematodes to evaluate the adverse effects of organic solvents on behavior. The speed of freely moving nematodes on an agar plate was measured, and the behavioral states were determined, namely forward movement, reverse, and pause, based on the speed and direction during locomotion (Figure 1A–C, Supporting Movie 1). First, the adverse effects of exposure to 10 kinds of monohydric alcohol were tested, whose carbon numbers are C1 to C7. The speed during forward movement and the ratio of the immobile state were quantified after exposure to the alcohols at several concentrations for 15 to 120 min (Figure 1D, Supporting Figure 1). For example, after exposure to 0.5% benzyl alcohol, the ratio of the immobile state and the speed of locomotion were substantially increased and decreased, respectively. The ratio of the immobile state increased to more than 0.5 after exposure for more than 1 h, and the speed decreased to less than 50% after exposure for more than 30 min (Figure 1D, Supporting Movie 2). The minimum concentrations of each alcohol that increased the immobility state to more than 0.5 or decreased the speed of locomotion to less than 50% after chemical exposure were determined, and the concentration was defined as MC_<50_. The MC_<50_ was established for the 10 monohydric alcohols (Figure 1D, Supporting Figure 1). It was reported that the inhibitory effects of alcohols on growth are increased with the increasing number of carbons of the alcohol in the yeast *Saccharomyces cerevisiae* [8]. The MC_<50_ for *C. elegans* locomotion was compared with the carbon numbers of the alcohols to which they were exposed. The MC_<50_ for immobility and locomotion speed exhibited a strong correlation with the carbon numbers of the alcohols (Figure 2A). These results suggest that the behavioral toxicity of monohydric alcohol can be determined by the immobility rate or locomotion speed and is proportional to its carbon number in *C. elegans*. To quantify the behavioral toxicity of chemicals, the locomotion speed was mainly used as an endpoint for subsequent analyses.

### Behavioral toxicity of organic solvents is proportional to the chemical’s lipid solubility

Next, the behavioral toxicity of organic solvents with distinct functional groups was determined, namely ketone, acetate ester, ether, cellosolve, and cellosolve acetate, using locomotion speed as an endpoint (Supporting Figure 2–3). Similar to the adverse effects of alcohol, exposure to these organic solvents decreased the locomotion speed on an agar plate (Supporting Figure 2–3). Furthermore, the toxicities of chemicals with ketone or cellosolve groups were proportional to the chemical’s carbon numbers (Supporting Figure 4). However, compared to the correlation of the MC_<50_ with the carbon number of the 10 kinds of alcohols (Figure 2A) that of the 30 kinds of organic solvents was low, presumably because the distinct functional groups exert various effects on locomotion (Figure 2B). The MC_<50_ was then compared with the lipid solubility of the organic solvents, which is an important property for several biological processes, such as skin permeability and the anesthetic effect. High levels of correlation were observed when the MC_<50_ was compared with the octanol–water partition coefficient, which reflects lipid solubility (Figure 2C, Supporting Figure 5). Meanwhile, no significant correlation was observed between the MC_<50_ and the chemical melting points, which affects skin permeability (Supporting Figure 6). These results suggest that lipid solubility is an important property for organic solvent toxicity of locomotion in *C. elegans*.

### Low molecular ketones and ethers cause body shrinkage followed by relaxation

Levamisole, an anthelmintic, has been known to cause paralysis in *C. elegans*. Levamisole acts as a potent agonist of acetylcholine receptors, and exposure to it causes muscle contraction followed by muscle relaxation [18]. It was confirmed that levamisole reduced the locomotion speed and eventually ceased the movement at 100 µM after 1 h of exposure. (Supporting Figure 7A, B). The impaired locomotion was associated with body shrinkage (Supporting Figure 7C, D, middle). Most nematodes were completely paralyzed by levamisole exposure at 1000 µM, and body relaxation was observed after 1 h of exposure (Supporting Figure 7A–C, D, right). Similar to these observations, exposure to a ketone, 2-butanone, or an ether, tetrahydrofuran, gradually decreased the body length at ranges of effective concentrations for impaired locomotion, which was followed by relaxation at concentrations that led to complete paralysis (Figure 3A, Supporting Figure 2A, C, and 8). Other C3–C6 ketones and C4 ethers, except 2-hexanone, had similar effects (Figure 3B). Alternatively, most alcohol, acetate ester, cellosolve, and cellosolve acetate groups of chemicals had no significant effect on body shrinkage except 2-hexanol and methyl acetate (Supporting Figure 9). These results imply that low molecular ketones and ethers promote muscle contraction followed by muscle relaxation similar to that of levamisole.

**Figure 7.**
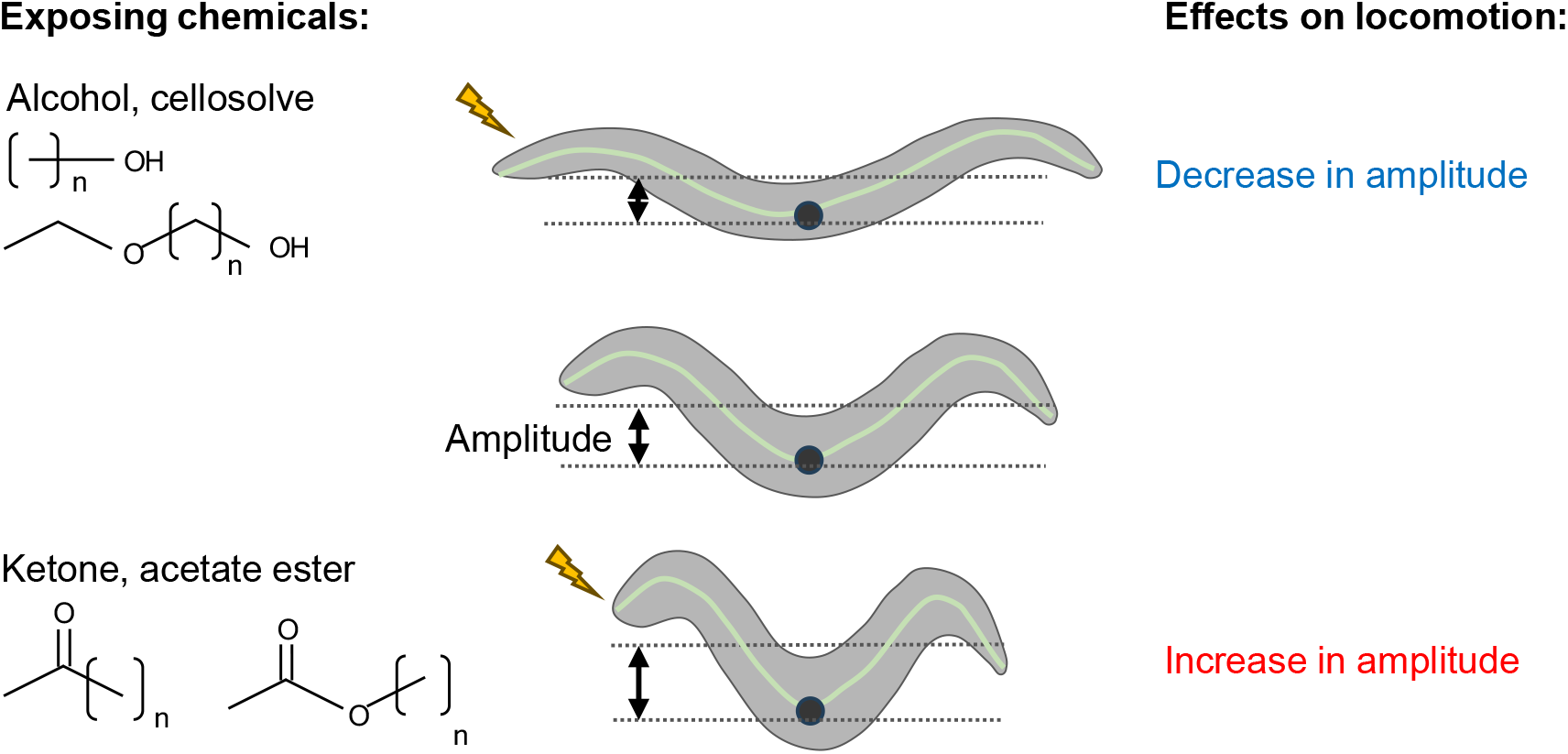
Summary of the adverse effects of organic solvents on the amplitude of body bending during locomotion. Organic solvents cause adverse effects on the amplitudes of body bending during locomotion in opposite ways depending on the chemical’s functional groups. Exposure to short-chain alcohols and cellosolve, which have a hydroxyl group, decreases the amplitude of body bending (top), whereas exposure to short-chain ketones and acetate esters, which have a ketone group, increases the amplitude of body bending (bottom).

To assess whether 2-butanone and tetrahydrofuran affect muscle functions via levamisole-sensitive acetylcholine receptors, the effects of exposure to these chemicals were examined in a loss-of-function mutant of UNC-29, which is an essential subunit of levamisole-sensitive acetylcholine receptors expressed in the body-wall muscle [19]. It was first confirmed that the loss-of-function *unc-29(e193)* mutant was resistant to levamisole exposure, which is consistent with a previous report that the acetylcholine receptors containing the UNC-29 subunit are the primary target of levamisole in the muscle (Supporting Figure 10A). In contrast, exposure to 2-butanone or tetrahydrofuran caused substantial locomotion impairment in the *unc-29* mutant (Supporting Figure 10B, C). Rather, the *unc-29* mutant was sensitive to organic solvents, including 2-butanone and tetrahydrofuran, when the immobility rate and locomotion speed were used as endpoints (Supporting Figure 11). These results suggest that low molecular ketones and ethers impair locomotion, at least in part, in a UNC-29-independent manner.

### Recovery from paralysis caused by organic solvent exposure

To examine whether locomotion impairment after organic solvent exposure was reversible, the locomotion of nematodes was assessed after recovery in a buffer without organic solvent. Nematodes were exposed to organic solvents for 2 h at concentrations that caused complete paralysis and then were soaked in a buffer without the solvents. The impaired locomotion was recovered within 2 h of soaking in a buffer after paralysis caused by exposure to most of the organic solvents except benzyl alcohol, 2-pentanone, 2-hexanone, and ethyl acetate (Figure 4). Complete paralysis after exposure to benzyl alcohol, 2-pentanone, 2-hexanone, and ethyl acetate for shorter periods, 15 or 30 min, was recovered by soaking in a buffer without the solvents (Supporting Figure 12). These results suggest that nematodes are paralyzed after short-term exposure to organic solvents, which is reversible, whereas long-time exposure to some organic solvents can cause irreversible locomotion impairment. These phenotypes are reminiscent of those after exposure to a human anesthetic, halothane, in *C. elegans* [20].

### Organic solvents have qualitatively different effects on the amplitude of body bends during locomotion

Apart from the general adverse effects of organic solvents on locomotion, is there a specific effect that is qualitatively different based on the solvent’s chemical characteristics, such as its functional group (Figure 5A)? It has been reported that ethanol exposure decreases the amplitude of body bends as well as speed during locomotion in *C. elegans* [21]; therefore, the amplitude of the body bends during locomotion was quantified (Figure 5B, C). As reported for ethanol exposure, exposure of alcohols, including methanol, isopropanol, and 2-butanol, for 15 min decreased the amplitude of body bends during locomotion at concentrations lower than MC_<50_ (Figure 5D, Supporting Movie 3). Similar effects on the body bends were observed by exposure to cellosolves, including methyl cellosolve, ethyl cellosolve, and isopropyl cellosolve and cellosolve acetate; these organic solvents decreased the amplitude of the body bends (Supporting Figure 13A, B).

Interestingly, when nematodes were exposed to ketone and acetate ester, the effects on the body bends were the opposite of those exposed to alcohols and cellosolves. Exposure to ketones, including acetone, 2-butanone, and 2-pentanone, and acetate esters, including methyl acetate, ethyl acetate, and isopropyl acetate, for 15 min significantly increased the amplitude of body bends during locomotion at the concentration lower than MC_<50_ (Figure 5E, Supporting Figure 13C, Supporting Movie 4). The adverse effects of ether were intricate: 1,4-dioxane decreased the amplitude of the body bend; diethyl ether did not affect the body bends; and tetrahydrofuran decreased and increased the amplitude of the body bends at concentrations of 0.5% and 2%, respectively. (Supporting Figure 13D, Supporting Movie 5). These results suggest that exposure to organic solvents affects the posture during locomotion differently at concentrations lower than those that cause cessation of motility. Taken together with the report that the amplitude of the body bends is regulated by rigorous functions of motor neurons that connect with the neck muscles, these results imply that organic solvents have distinct effects on the motor neurons that control the body bends.

### Organic solvents lead to deficits in sensory processing at concentrations lower than those that affect locomotion

To explore whether organic solvent exposure affects sensory processing in *C. elegans*, salt chemotaxis learning, a paradigm of behavioral plasticity, were examined [16]. *C. elegans* responds to various environmental chemicals, including NaCl, and exhibits behavioral plasticity appropriate to the situation [22]. The nematodes are attracted toward NaCl, whereas after prolonged exposure to high concentrations of NaCl under starvation conditions, they avoid NaCl. This behavioral plasticity termed “salt chemotaxis learning” is interpreted as a kind of learned behavior where nematodes memorize NaCl concentrations in their environment experienced with starvation and avoid the environment without food using the NaCl concentration as a repulsive cue [23]. Organic solvents were added to a buffer for conditioning, in which the nematodes were exposed to the organic solvents in the presence or absence of NaCl for 1 h to evaluate the effects of the organic solvents on NaCl avoidance or attraction, respectively (Supporting Figures 14). Organic solvents were added at concentrations that had no significant effect on the locomotion speed on an agar plate (Supporting Figures 1–3). A significant defect was observed in NaCl attraction after exposure in the absence of NaCl when ethyl acetate was applied. Meanwhile, significant defects were observed after exposure in the presence of NaCl when exposed to any of the organic solvents tested, namely alcohol, ketone, acetate ester, ether, and cellosolve, except methyl cellosolve (Figure 6A). These results suggest that these organic solvents cause defects in salt chemotaxis learning at concentrations lower than those that affect locomotion speed. Substantial defects in salt chemotaxis learning were not observed by exposure to any of the organic solvents for 1 min after starvation conditioning in the presence of NaCl, which suggests that prolonged exposure to organic solvents during conditioning is required for the establishment of adverse effects on salt chemotaxis learning (Figure 6B).

Glutamate is a neurotransmitter that regulates the functions of sensory neural circuits in chemotaxis learning in *C. elegans* [24]. Indeed, mutants of *eat-4*, which encodes a glutamate transporter, exhibited reduced attraction and avoidance of NaCl in salt chemotaxis learning (Figure 6C). Organic solvent exposure had no significant effect on salt chemotaxis learning in nematodes with an *eat-4* mutant background, which is consistent with the notion that the organic solvents affect the sensory neural circuit that functions in salt chemotaxis learning (Figure 6C). Insulin signaling plays a pivotal role in the regulation of salt chemotaxis learning. DAF-2c, which encodes an insulin receptor isoform localized in the neuronal axon, regulates the neuronal plasticity of the NaCl-sensing sensory neuron ASER, and a *daf-2c* mutant exhibits substantial defects in salt chemotaxis learning [25] (Figure 6D). Organic solvent exposure promoted the defect in NaCl avoidance or caused defects in attraction in nematodes with a *daf-2c* mutant background, suggesting that organic solvent exposure adversely affects sensory processing, at least in part, independent of DAF-2c signaling in salt chemotaxis learning (Figure 6D).

## Discussion

To reduce the health hazards of industrial chemicals, which are continuously increasing, it is important to fundamentally understand the toxic effects of chemicals on organisms. Because of its fast lifecycle and proliferative capacity, *C. elegans* is useful for high-throughput screening, including toxic chemical screening [10, 11, 26]. In this study, a high-throughput method was developed to assess the acute toxicity of organic solvents. The nematodes were exposed to chemicals by soaking in a buffer and were transferred onto an agar plate to capture a 1-min movie. The behaviors of the nematodes were then analyzed using a commercially available tracking software, WormLab. This method is simple, and no special skills are required. Using this method, the behavioral toxicity of 30 organic solvents commonly used in industries was comprehensively analyzed. Exposure to any of the organic solvents gradually decreased the movement, and locomotion eventually ceased. Because the paralyzed nematodes were recovered by soaking in a buffer without organic solvents, the locomotion impairments were reversible. When the locomotion speed was used as an endpoint, the extent of the behavioral toxicities was proportional to the chemical’s lipid solubility. This relationship was reminiscent of that in humans called the Meyer–Overton relationship, in which there is a positive relationship between the anesthetic effects and the lipid solubilities of inhalation anesthetics, such as halothane and isoflurane [7]. It has been reported that there is a positive relationship between volatile anesthetics and impaired locomotion in *C. elegans* [13]. Therefore, this study confirmed the availability of assessing the general anesthetic effects of industrial chemicals using *C. elegans*.

Exposure to organic solvents, such as short-chain ketones and ethers, caused body shrinkage followed by muscle relaxation, which is similar to that of levamisole, an anthelmintic agent. Levamisole is a well-known potent agonist of acetylcholine receptors, including UNC-29, in *C. elegans* [18]. As previously reported, the *unc-29* mutant was resistant to locomotion impairment caused by levamisole exposure. Conversely, the *unc-29* mutant was sensitive to organic solvents, including alcohol, ketone, ether, and acetate ester, and required lower concentrations of the organic solvents to induce locomotion impairment than those in the wild type. Acetylcholine signaling plays an important role in the neuromuscular junctions during locomotion [18]. Ketone and ether exposure may affect acetylcholine signaling in the neuromuscular junction as well as multiple functions of neurons and/or muscles. Because exposure to alcohol and cellosolve did not cause significant body shrinkage, the contraction effect of muscle may be specific to certain types of organic solvents, including ketone and ether.

In addition to the general effect of organic solvents on behavior, it was found that organic solvents have specific effects on behavioral patterns; they have different qualitative effects on the amplitude during undulatory locomotion based on the difference of the chemical’s functional groups. Alcohols and cellosolves, which have a hydroxyl group, and ketones and acetate esters, which have a ketone group, decrease and increase the body bend amplitude during locomotion, respectively (Figure 7). It is possible that a hydroxyl group and a ketone group may affect the motor neural circuits that regulate the amplitude of head bending in different manners. The amplitude of head bending during undulatory locomotion is regulated by several motor neurons that innervate the head and neck muscles. The cholinergic SMB motor neurons negatively regulate the amplitude of head bending, as SMB ablation causes a strong increase in the amplitude of head bending [14]. Conversely, the cholinergic SMD motor neurons positively regulate the amplitude of head bending. The GABAergic RMD motor neurons negatively regulate the amplitude of head bending via SMD inhibition [15]. Organic solvents with a hydroxyl group and a ketone group may have different effects on the same motor neurons to induce opposite effects on the amplitude. Alternatively, those organic solvents may have an effect on different types of motor neurons with opposite functions. Further studies are required to elucidate the molecular and cellular mechanisms of the different qualitative adverse effects of organic solvents on behavioral patterns and their possible conservation across species.

Organic solvents also had adverse effects on learned behavior termed “salt chemotaxis learning” at concentrations lower than those that affect locomotion speed or the amplitude of head bending. Expression of these adverse effects requires glutamatergic signaling, which plays a pivotal role in the control of the sensory neural circuits. This suggests that organic solvents have adverse effects on the sensory processing functions at lower concentrations compared to those of motor neural circuits [27] (Supporting Figure 15). Nematodes respond to various ambient chemicals, including volatile and water-soluble chemicals. Water-soluble chemicals, including NaCl, are mainly received by sensory neurons called ASE. The ASE neurons extend neuronal processes to the nose tip of the head, and the ciliated ends are exposed to the outside environment through the sensory organ called the amphid [28]. There are molecular mechanisms that regulate sensory transduction and sensory processing in the ciliated ends of ASE [29]. Organic solvents may directly affect the mechanisms for sensory transduction and processing in the ASE cilia, which are directly exposed to the environment. This may be a reason for the high susceptibility of the sensory processing system compared to motor neurons, whose neuronal processes are located beneath the hypodermis in *C. elegans*. Alternatively, compared to the motor control of the robust rhythmic movement, the sensory processing system may be more sensitive to respond to sensory information, such as subtle changes in the environment and multiple sensory stimuli. Therefore, the sensory processing system may be more susceptible to organic solvent exposure than the motor control systems. It is important to understand the cellular and molecular mechanisms that underlie the adverse effects of organic solvent exposure in future studies to elucidate the mechanisms of chemical toxicities on organisms, which can then be applied to toxicity assessment for other organisms, including humans.

## Acknowledgments

I would like to thank Drs. Tatsushi Toyooka, Rui-Sheng Wang, and Makiko Nakano at the National Institute of Occupational Safety and Health, Japan, for their helpful discussions and support. The *C. elegans* and *E. coli* strains were provided by Dr. Yuichi Iino’s lab and the *Caenorhabditis* Genetics Center.

## References

1. Ichimata S, Hata Y, Zaimoku R, Nishida N. Acute benzyl alcohol intoxication: An autopsy case report. Medicine (Baltimore). 2023;102(13):e33395. doi: 10.1097/MD.0000000000033395. PubMed PMID: 37000071; PubMed Central PMCID: PMCPMC10063254.

2. Inada M, Kato H, Maruyama H, Okada E, Imai F, Inada S. Toxic benzyl alcohol inhalation: Altered mental status with metabolic acidosis and hyperammonemia. Am J Emerg Med. 2022;57:234.e3-.e5. Epub 20220411. doi: 10.1016/j.ajem.2022.04.005. PubMed PMID: 35466010.

3. Winder C, Azzi R, Wagner D. The development of the globally harmonized system (GHS) of classification and labelling of hazardous chemicals. J Hazard Mater. 2005;125(1-3):29–44. doi: 10.1016/j.jhazmat.2005.05.035. PubMed PMID: 16039045.

4. Jonai H. Implementation of the GHS in Japan. Ind Health. 2008;46(5):443–7. doi: 10.2486/indhealth.46.443. PubMed PMID: 18840933.

5. Meyer HH. Zur Theorie der alkoholnarkose. I. Mitt. Welche eigenschaft der anasthetika bedingt ihre narkotische wirkung. Arch Exp Pathol Pharmak. 1899;42:109–19.

6. Overton E. Studien uber die Narkose, Zugleich ein Beitrag zur allgemeinen Pharmakologie. Jena: Gustav Fischer. 1901.

7. Kopp Lugli A, Yost CS, Kindler CH. Anaesthetic mechanisms: update on the challenge of unravelling the mystery of anaesthesia. Eur J Anaesthesiol. 2009;26(10):807–20. doi: 10.1097/EJA.0b013e32832d6b0f. PubMed PMID: 19494779; PubMed Central PMCID: PMCPMC2778226.

8. Matsumoto A, Uesono Y. Physicochemical Solubility of and Biological Sensitivity to Long-Chain Alcohols Determine the Cutoff Chain Length in Biological Activity. Mol Pharmacol. 2018;94(6):1312–20. Epub 20181005. doi: 10.1124/mol.118.112656. PubMed PMID: 30291172.

9. Hartman JH, Widmayer SJ, Bergemann CM, King DE, Morton KS, Romersi RF, et al. Xenobiotic metabolism and transport in. J Toxicol Environ Health B Crit Rev. 2021;24(2):51–94. Epub 20210222. doi: 10.1080/10937404.2021.1884921. PubMed PMID: 33616007; PubMed Central PMCID: PMCPMC7958427.

10. Hunt PR. The C. elegans model in toxicity testing. J Appl Toxicol. 2017;37(1):50–9. Epub 20160722. doi: 10.1002/jat.3357. PubMed PMID: 27443595; PubMed Central PMCID: PMCPMC5132335.

11. Boyd WA, Smith MV, Co CA, Pirone JR, Rice JR, Shockley KR, et al. Developmental Effects of the ToxCast™ Phase I and Phase II Chemicals in Caenorhabditis elegans and Corresponding Responses in Zebrafish, Rats, and Rabbits. Environ Health Perspect. 2016;124(5):586–93. Epub 20151023. doi: 10.1289/ehp.1409645. PubMed PMID: 26496690; PubMed Central PMCID: PMCPMC4858399.

12. International Organization for Standardization. Water and soil quality — Determination of the toxic effect of sediment and soil samples on growth, fertility and reproduction of Caenorhabditis elegans (Nematoda). 2020;ISO 10872:2020.

13. Morgan PG, Cascorbi HF. Effect of anesthetics and a convulsant on normal and mutant Caenorhabditis elegans. Anesthesiology. 1985;62(6):738–44. doi: 10.1097/00000542-198506000-00007. PubMed PMID: 4003794.

14. Gray JM, Hill JJ, Bargmann CI. A circuit for navigation in Caenorhabditis elegans. Proc Natl Acad Sci U S A. 2005;102(9):3184–91. Epub 20050202. doi: 10.1073/pnas.0409009101. PubMed PMID: 15689400; PubMed Central PMCID: PMCPMC546636.

15. Shen Y, Wen Q, Liu H, Zhong C, Qin Y, Harris G, et al. An extrasynaptic GABAergic signal modulates a pattern of forward movement in Caenorhabditis elegans. Elife. 2016;5. Epub 20160503. doi: 10.7554/eLife.14197. PubMed PMID: 27138642; PubMed Central PMCID: PMCPMC4854516.

16. Tomioka M, Adachi T, Suzuki H, Kunitomo H, Schafer WR, Iino Y. The insulin/PI 3-kinase pathway regulates salt chemotaxis learning in Caenorhabditis elegans. Neuron. 2006;51(5):613–25. doi: 10.1016/j.neuron.2006.07.024. PubMed PMID: 16950159.

17. Brenner S. The genetics of Caenorhabditis elegans. Genetics. 1974;77(1):71–94. doi: 10.1093/genetics/77.1.71. PubMed PMID: 4366476; PubMed Central PMCID: PMCPMC1213120.

18. Rand JB. Acetylcholine. WormBook. 2007:1–21. Epub 20070130. doi: 10.1895/wormbook.1.131.1. PubMed PMID: 18050502; PubMed Central PMCID: PMCPMC4781110.

19. Hunt PR, Welch B, Camacho J, Bushana PN, Rand H, Sprando RL, et al. The worm Adult Activity Test (wAAT): A de novo mathematical model for detecting acute chemical effects in Caenorhabditis elegans. J Appl Toxicol. 2023;43(12):1899–915. Epub 20230808. doi: 10.1002/jat.4525. PubMed PMID: 37551865.

20. Morgan PG, Kayser EB, Sedensky MM. C. elegans and volatile anesthetics. WormBook. 2007:1–11. Epub 20070503. doi: 10.1895/wormbook.1.140.1. PubMed PMID: 18050492; PubMed Central PMCID: PMCPMC4781344.

21. McIntire SL. Ethanol. WormBook. 2010:1–6. Epub 20100429. doi: 10.1895/wormbook.1.40.1. PubMed PMID: 20432508; PubMed Central PMCID: PMCPMC4781098.

22. Zhang Y, Iino Y, Schafer WR. Behavioral plasticity. Genetics. 2024;228(1). doi: 10.1093/genetics/iyae105. PubMed PMID: 39158469.

23. Kunitomo H, Sato H, Iwata R, Satoh Y, Ohno H, Yamada K, et al. Concentration memory-dependent synaptic plasticity of a taste circuit regulates salt concentration chemotaxis in Caenorhabditis elegans. Nat Commun. 2013;4:2210. doi: 10.1038/ncomms3210. PubMed PMID: 23887678.

24. Cheng D, Lee JS, Brown M, Ebert MS, McGrath PT, Tomioka M, et al. Insulin/IGF signaling regulates presynaptic glutamate release in aversive olfactory learning. Cell Rep. 2022;41(8):111685. doi: 10.1016/j.celrep.2022.111685. PubMed PMID: 36417877.

25. Tomioka M, Jang MS, Iino Y. DAF-2c signaling promotes taste avoidance after starvation in Caenorhabditis elegans by controlling distinct phospholipase C isozymes. Commun Biol. 2022;5(1):30. Epub 20220111. doi: 10.1038/s42003-021-02956-8. PubMed PMID: 35017611; PubMed Central PMCID: PMCPMC8752840.

26. Shin N, Cuenca L, Karthikraj R, Kannan K, Colaiácovo MP. Assessing effects of germline exposure to environmental toxicants by high-throughput screening in C. elegans. PLoS Genet. 2019;15(2):e1007975. Epub 20190214. doi: 10.1371/journal.pgen.1007975. PubMed PMID: 30763314; PubMed Central PMCID: PMCPMC6375566.

27. Sato H, Kunitomo H, Fei X, Hashimoto K, Iino Y. Glutamate signaling from a single sensory neuron mediates experience-dependent bidirectional behavior in Caenorhabditis elegans. Cell Rep. 2021;35(8):109177. doi: 10.1016/j.celrep.2021.109177. PubMed PMID: 34038738.

28. Bargmann CI. Chemosensation in C. elegans. WormBook. 2006:1–29. Epub 20061025. doi: 10.1895/wormbook.1.123.1. PubMed PMID: 18050433; PubMed Central PMCID: PMCPMC4781564.

29. Ferkey DM, Sengupta P, L’Etoile ND. Chemosensory signal transduction in Caenorhabditis elegans. Genetics. 2021;217(3). doi: 10.1093/genetics/iyab004. PubMed PMID: 33693646; PubMed Central PMCID: PMCPMC8045692.

